# The microbiota influences the *Drosophila melanogaster* life history strategy

**DOI:** 10.1101/471540

**Authors:** Amber W. Walters, Melinda K. Matthews, Rachel Hughes, Jaanna Malcolm, Seth Rudman, Peter D. Newell, Angela E. Douglas, Paul S. Schmidt, John M. Chaston

## Abstract

**Abstract:** Organismal life history traits are ideally adapted to local environments when an organism has a fitness advantage in one location relative to conspecifics from other geographies. Local adaptation has been best studied across, for example, latitudinal gradients, where organisms may tradeoff between investment in traits that maximize one, but not both, fitness components of somatic maintenance or reproductive output in the context of finite environmental resources. Latitudinal gradients in life history strategies are traditionally attributed to environmentally mediated selection on an animal’s genotype, without any consideration of the possible impact of associated microorganisms (‘microbiota’) on life history traits. Here we show that in *Drosophila melanogaster*, a key organism for studying local adaptation and life history strategies, associated microorganisms can drive life history variation. First, we reveal that an isogenic fly line reared with different bacteria vary the investment in early reproduction versus somatic maintenance, with little resultant variation in lifetime fitness. Next, we show that in wild *Drosophila* the abundance of these same bacteria was correlated with the latitude and life history strategy of the flies, and bacterial abundance was driven at least in part by host genetic selection. Finally, by eliminating or manipulating the microbiota of fly lines collected across a latitudinal gradient, we reveal that host genotype contributes to latitude-specific life history traits independent of the microbiota; but that the microbiota can override these host genetic adaptations. Taken together, these findings establish the microbiota as an essential consideration in local adaptation and life history evolution.

**Significance statement:** Explanations of local adaptation have historically focused on how animal genotypes respond to environmental selection. Although the impact of variation in host life histories on the composition of the microbiota has been investigated for many associations, the scale and pattern of microbial effects on host life history strategy are largely unknown. Here we demonstrate in the fruit fly *Drosophila melanogaster* that microbiota effects on host life history strategy in the laboratory are matched by patterns of microbiota composition in wild host populations. In particular, microbiota composition varies with latitude and the effects of the microbiota on life history traits are greater than host genetic adaptations. Together, these findings demonstrate that the microbiota plays an important role in local adaptation.

## Introduction

Life history tradeoffs have long been recognized as a widespread feature of local adaptation and have been the focus of many empirical studies (1, 2). An animal’s life history reflects its allocation of resources and time to maximize reproductive output, subject to natural selection and tradeoffs along a ‘fast-slow’ continuum (3-5). At the ‘fast’ end, organisms develop to reproductive maturity more quickly and have high early fecundity; whereas a ‘slow’ lifestyle favors somatic maintenance and lower initial reproduction across longer lifespan (6, 7). These insights have been developed in the context of an environment-genotype centric framework, focused on geography-specific environmental selection on the organismal genotype mediated, for example, by temperature or photoperiod (8, 9). The consequent variation in genotype has been linked to various physiological and behavioral characters collectively described as the pace of life syndrome (4, 10).

The rationale for this study is the abundant evidence that key life history traits (e.g. development rate, fecundity, lifespan) and correlated physiological traits can be influenced by the presence and composition of associated microorganisms (‘microbiota’) (11-17). To date, research on interactions between microbiota and the life history strategy of the host has focused exclusively on how inter- and intra-specific variation in host life history strategies influence the microbiota, e.g. (18, 19). The reverse question –the impact of the microbiota on the life history strategy of the host, including the evolution of locally-adapted populations – has, to our knowledge rarely been considered (20), and has not been investigated empirically.

*Drosophila melanogaster* is an excellent system to address the impact of the microbiota of host life history strategy because both its life history and microbiota are well-studied. Considering its life history first, tradeoffs and their role in local adaptation have been demonstrated, especially in relation to latitudinal clines in allele frequencies for fitness-associated traits (21-25), candidate genes (26-30), and genome-wide patterns (31-33). In particular, *D. melanogaster* adopt different life history strategies across a latitudinal gradient in the eastern United States. Flies at high latitudes, e.g. Maine, occupy the ‘slower’, somatic maintenance-promoting end of the fast-slow continuum (long lifespans and stress survival, high fat storage); whereas flies at low latitudes, e.g. Florida, invest in rapid development and early reproduction (21, 34, 35). Turning to the gut microbiota, a growing body of research has revealed that the gut microbiota of *D. melanogaster* is of low diversity, represented by <100 species, usually dominated by acetic acid bacteria (AAB) of the family Acetobacteraceae, including *Acetobacter* species, or lactic acid bacteria (LAB) from the order Lactobacillales, including the genera *Lactobacillus*, *Enterococcus*, and *Leuconostoc* (36-41). As in many other animals, the *D. melanogaster* microbiota varies both among individual hosts and over time within an individual animal (42, 43), and this variation is shaped by both deterministic factors, e.g. host genotype, among-microbe interactions, diet composition (44-49) and stochastic processes of passive dispersal and ecological drift (50-53). The gut microbiota of *D. melanogaster* microbiota is also readily manipulated in the laboratory: it can be eliminated by bleach treatment; the dominant taxa are fully culturable; and microbial communities of defined composition can be administered by direct inoculation to bleach-sterilized fly eggs on a sterile diet, generating gnotobiotic flies (54). If no bacteria are reapplied, the resultant “axenic” insects develop and reproduce with no evidence of generalized malaise (55).

The basis for this study is the observation that presence and composition of the *D. melanogaster* microbiota affect key traits of *D. melanogaster* that underpin life history strategy, including development rate, lifespan and fecundity (55-63). We hypothesized that the microbiota might, therefore, influence patterns of local adaptation in *D. melanogaster*. We asked three questions: 1) How does the microbiota influence traits contributing to the life history strategy of their host? 2) Does the taxonomic composition of the microbiota in *D. melanogaster* vary with geographical location along the latitudinal cline in eastern USA? 3) What are the relative contributions of host genotype and the microbiota in shaping local adaptation of the host along this cline? Using studies of both laboratory and wild populations of *D. melanogaster*, we reveal that (i) the identity of associated microorganisms influences the position of the flies along the fast-slow axis; (ii) relative abundances of key members of the microbiota in wild-caught flies correlate with life history traits and can be determined by host genetic selection; (iii), local adaptation of the host genotype is independent of the microbiota, but can be masked by microbiota effects. Together, these findings suggest that microbes are an essential consideration in evaluating the causal basis for local adaptation in their animal hosts.

## Results

### The microbiota influences D. melanogaster life history strategy

In an evaluation of previously collected datasets (59, 60, 64), we noticed correlated influences of Acetic Acid Bacteria (AABs) and Lactic Acid Bacteria (LABs) on *D. melanogaster* life history traits. Specifically, the isogenic *D. melanogaster* CantonS line tended to display faster development rates, higher feeding rates, lower lipid (TAG) levels, and lower starvation resistance when monoassociated with AABs than LABs, and these correlated relationships were statistically significant (Fig. 1). The correlations were specific to the investigated traits since starvation resistance and development and feeding rates were not correlated with glucose content (Fig. 1), a nutritional index that is not usually considered with other life history traits (65). Since the trait correlations were consistent with established patterns of life history tradeoffs in *D. melanogaster* (65, 66), we hypothesized that variation in microbial colonization could influence the position that individual flies occupy along the fast-slow continuum. To test this hypothesis, we associated the same fly line with many different bacterial strains, and quantified additional life history traits in *D. melanogaster*: lifespan, and fecundity. Lifespan was positively correlated with SR and TAG content; but negatively correlated with ‘fast’ traits (fecundity, development rate, and feeding rate, which were positively correlated); and uncorrelated with glucose content. Among all tested life history traits, the correlation coefficients were consistent with the predicted variation along the fast-slow axis, although two correlations with relatively low replication (n=12, n=14) were not significant (Fig. 1).

**Figure 1.**
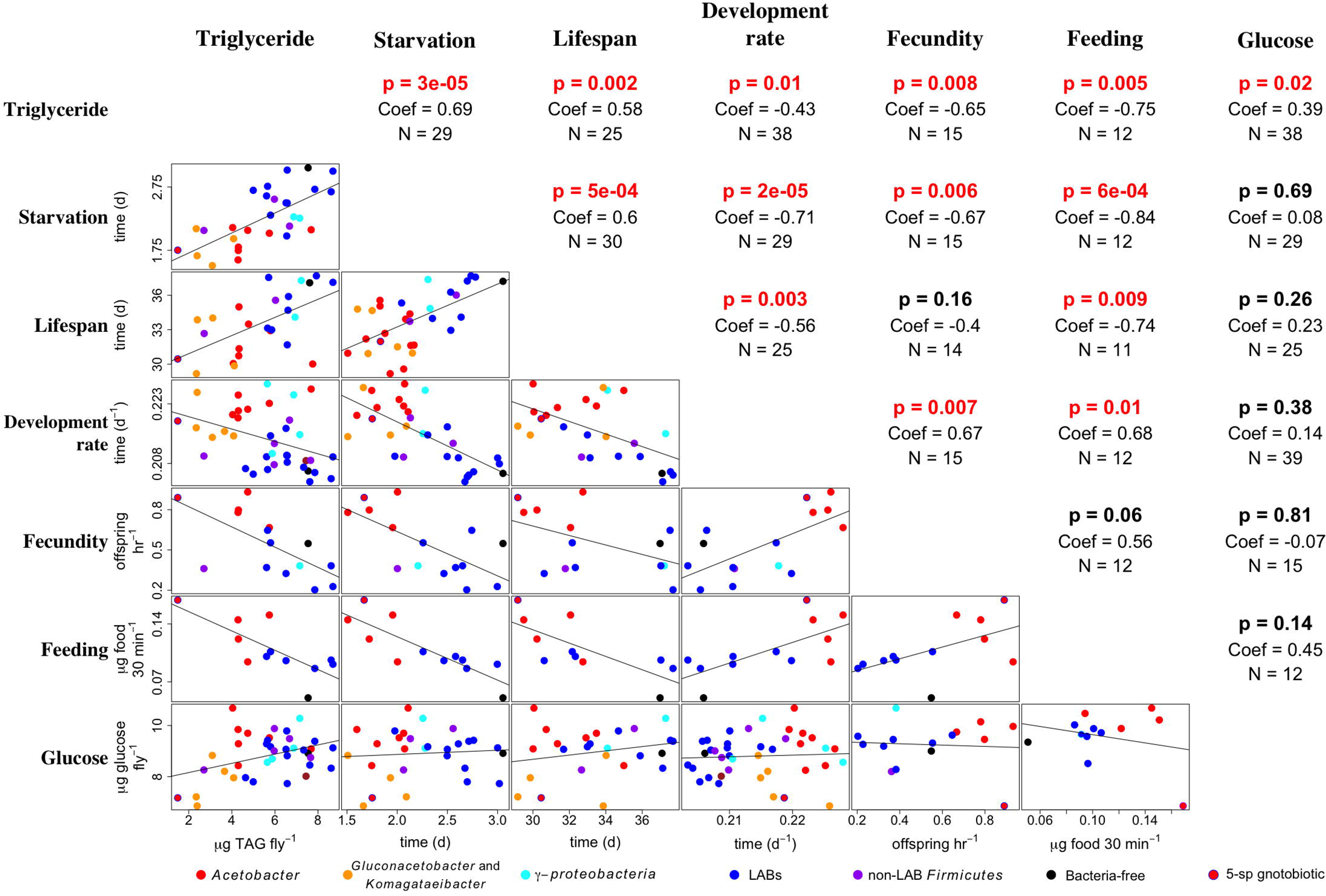
Microbial variation influences life history patterns in a laboratory reared isogenic fly line. Six life history traits were measured in *D. melanogaster* that were monoassociated with different bacterial species and reared on a YG diet: whole-body triacylglyceride content (Triglyceride), survival under starvation conditions (Starvation), lifespan, the rate of development to pupariation (Development rate), number of pupariating offspring produced in the first 2-4 days after eclosion (Fecundity), and feeding rate (Feeding). Fly whole body glucose content (Glucose), a trait that is not correlated with most other life history traits, was also measured. Mean trait values conferred by different bacteria are plotted in the bottom half of the table. The top half of the table shows the p-values (p), correlation coefficients (Coef), and number of different monoassociations (N). Each monoassociation usually had triplicate measures in three separate experiments. P-values that were significant after a Benjamini-Hochberg correction are shown in red. The data for triglyceride content, starvation resistance, development rate, and feeding rate were published previously (59, 60, 64).

### The microbiota has a greater impact on the temporal pattern of D. melanogaster fecundity than lifetime fitness

To test how the varied influences of different bacteria on fecundity and lifespan impact *D. melanogaster* fitness in the laboratory, we examined a time course of fecundity and longevity data in a matched set of female flies colonized with different bacteria (54). In the first of two experiments, we used 5 bacterial species isolated from the guts of the same *D. melanogaster* strain as used in Fig. 1 experiments (67) and administered either as single species or as a 5-species inoculum, with axenic insects as a control. Total viable offspring per hour did not vary significantly among the treatments (Fig. 2A), but microbial colonization dramatically affected early and late *D. melanogaster* fecundity (Fig 2B-C). These effects could not be attributed exclusively to differences in fly development since the average difference in fly development was < 12 hours (59). To quantify the fitness consequences of the among-treatment variation in fecundity and lifespan, we calculated the Eigenvalue lambda from Leslie matrices, using the data for each vial (68, 69). The only significant fitness difference was between flies reared with *L. fructivorans* or a 5-species community; fly fitness did not vary significantly among any other treatments (Fig. 2D).

**Figure 2.**
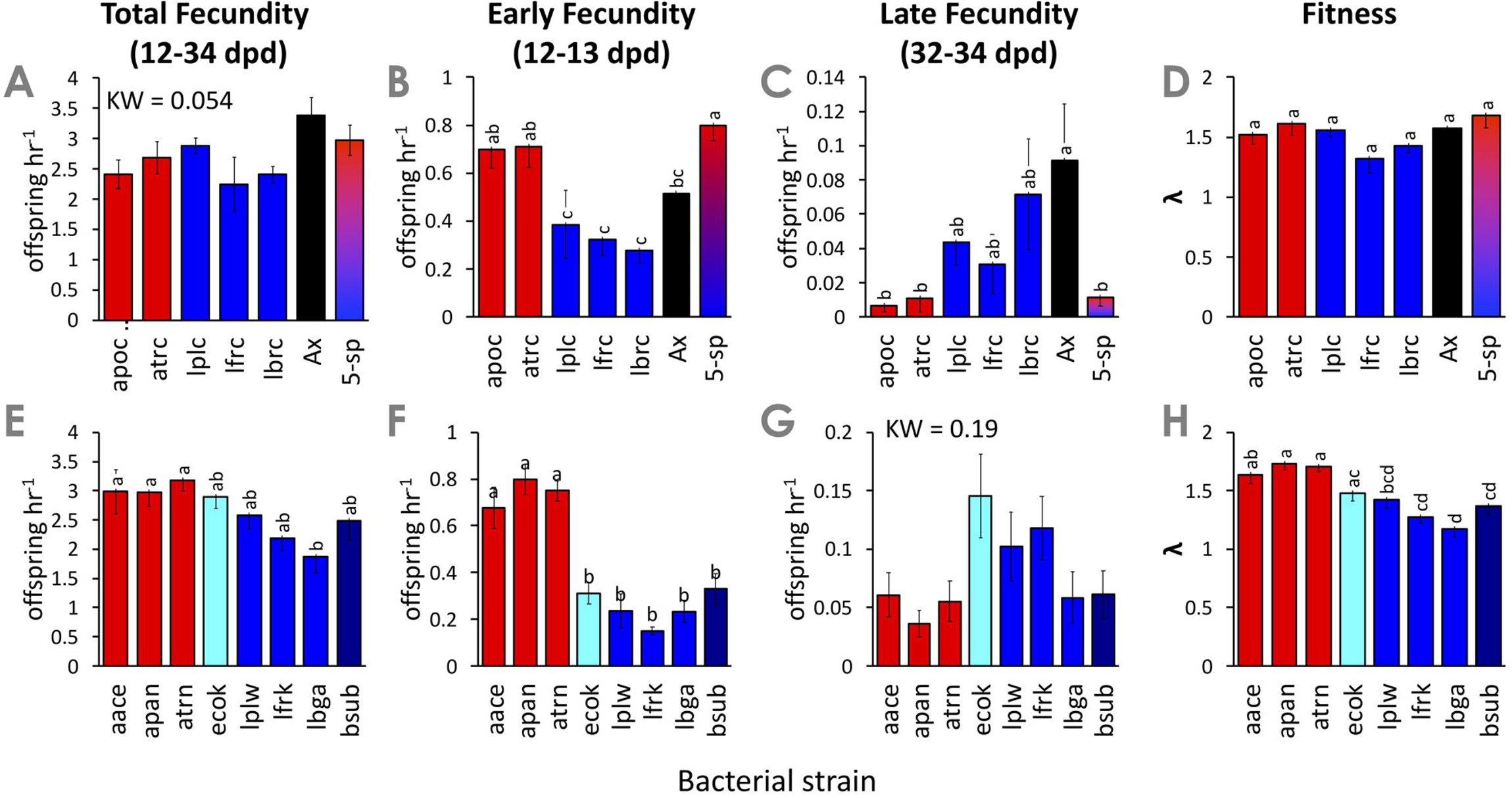
Microbial variation influences *D. melanogaster* fecundity, but not fitness. A-C)Pupariating offspring per hour produced by female *D. melanogaster* that were monoassociated with different bacterial species (y-axis). Aggregate intervals were A,E) 12-34 days post egg deposition (dpd), B,F) 12-13 dpd, or C,G) 32-34 dpd. D,H) Mean fitness lambda, calculated as the eigenvalue of a Leslie matrix constructed from the fecundity data in A-C or E-G and lifespan data collected for the same flies. Bar colors represent taxonomic assignments of the strains: *Acetobacter*, red; γ-proteobacteria, cyan; LABs, blue; non-LAB Firmicutes, purple; bacteria-free, black; 5-species gnotobiotic, red-blue gradient. N = 9 samples per treatment (triplicate vials in three separate experiments) except where vials were discarded for contamination or for early fecundity measures (N=6, two experiments, triplicate vials). KW = Kruskal-Wallis test p-value and corresponding p-value. If KW p > 0.05, no post-hoc test was performed. Otherwise, different letters over the bars represent statistically significant differences by a linear mixed model and Tukey post-hoc test. X-axis abbreviations are described in Table S4.

The second experiment tested if the fitness influences were limited to bacteria isolated form laboratory flies by quantifying the fecundity, lifespan and fitness of *D. melanogaster* individually associated with additional bacterial strains (Fig. 1). Similar trends to the first experiment were obtained, but with different patterns of significance. As in the first experiments, the *Acetobacter* species conferred higher early fecundity than the other taxa tested, and, although the effects on late fecundity were not statistically significant, a fitness differential between *Acetobacter* and the other bacteria was obtained (Fig. 2 E-H). The common feature of the two complementary experimental designs is that microorganisms have a stronger impact on the timing of fecundity, a key life history trait, than on lifetime fitness. Thus, the microbe-dependent differences in individual traits can lead to variation in the fast-slow strategy without reducing or promoting fitness.

### Host genetic selection influences latitudinal variation in D. melanogaster -associated LABs

To investigate the relevance of our laboratory studies to natural populations of *D. melanogaster*, we turned to the well-studied latitudinal cline in the eastern United States, where low latitude flies invest more in early reproduction than high-latitude flies (21, 35). We predicted that *D. melanogaster* from low latitude populations would bear *Acetobacter* and related bacterial taxa (AABs), while *Lactobacillus* and related taxa (LABs) would be associated with populations from higher latitudes. Using 16S rRNA marker gene sequencing we determined the relative abundance of AABs and LABs in wild flies from five latitudes along the eastern United States coast in the fall of 2009 (Tables S1-2). Reads were clustered at the order levels since LABs are an order level designation; AABs, from the family Acetobacteraceae represented 99.97% of the Rhodospirillales reads (data not shown), and Rhodospirillales reads are referred to as AABs hereafter for simplicity. Consistent with our predictions, relative AAB and LAB abundances were negatively and positively correlated with latitude, respectively (Fig. 3A). The other taxa tested did not vary significantly with latitude.

**Figure 3.**
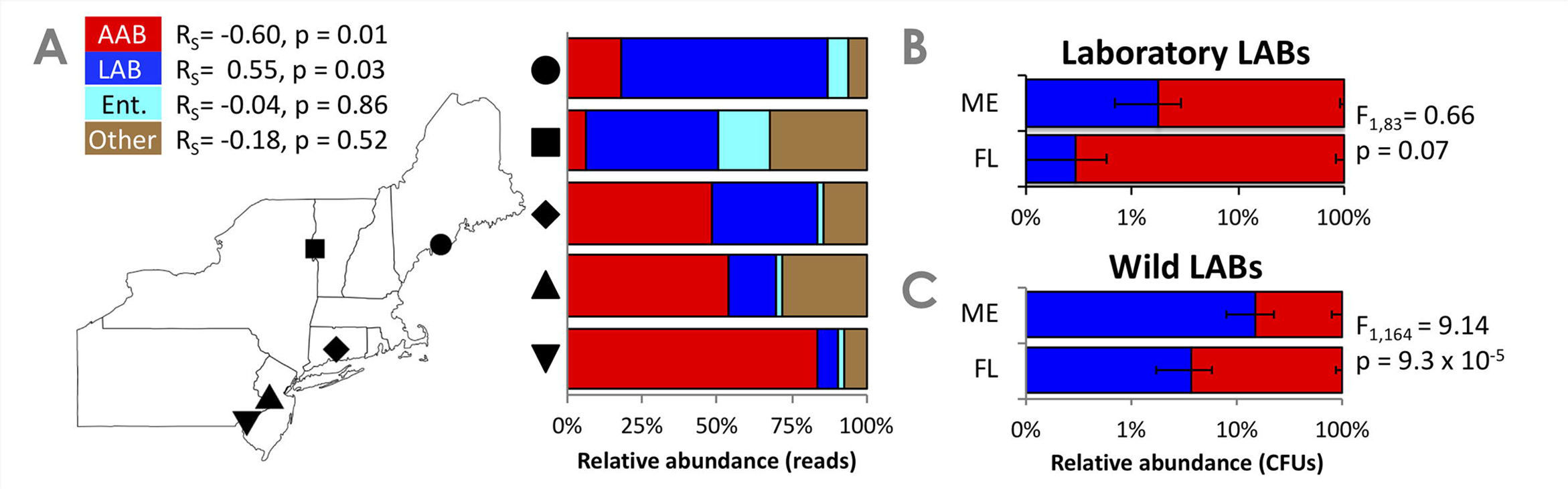
Latitudinal variation in the *D. melanogaster* microbiota. (A) Relative abundances of reads assigned to different bacterial orders in a 16S rRNA marker gene survey of *D. melanogaster* collected in 2009. Spearman’s rank correlations revealed significant positive and negative correlations between latitude and AAB or LAB read abundances, respectively. RS, Spearman’s rho. p, p-value. N=2-3 replicate pools of 10 flies each per geographic site. (B-C) Relative abundance of AABs (red) and LABs (blue) in isofemale lines derived from Maine (ME) – and Florida (FL)-wild populations, when reared under gnotobiotic conditions. B) Flies were reared with a 5-species microbiota, including 3 LABs isolated from laboratory *D. melanogaster*. C) Flies were reared with a 6-species gnotobiotic microbiota, including 4 LABs isolated from wild *D. melanogaster* The difference between relative LAB and AAB abundance was determined by a generalized linear mixed (GLM) effects model using a binomial family. F, F statistic of the GLM.

We then tested if host genetic selection on the microbiota could contribute to the observed latitude-specific microbiota differences. We inoculated bacteria-free fly lines that were recently derived from wild fly populations collected at the extrema of the eastern US (Maine, ME = high latitude, Florida, FL = low latitude) with defined microbial communities of AABs and LABs. The first experiment using our standard 5-species bacterial community derived from laboratory *Drosophila* (as in Fig. 2A-D) obtained a nearly significant difference in the relative abundance of AABs and LABs between the ME- and FL genotype flies (p= 0.07), and low overall relative abundance of LABs (Fig. 3B). The second experiment replaced the laboratory LABs by 4 LABs isolated from wild *Drosophila* (Table S3), yielding a four-fold increase of LABs in ME flies, relative to the first experiment, and a statistically significant difference between relative LAB abundance in ME and FL flies (Fig. 3C). These differences were largely driven by LAB abundance in the ME67 line and supported by the lack of any detectable LABs (limit of detection = 20 CFU fly^-1^) in 3 of 4 FL-derived fly lines (Fig. S2). The absolute AAB abundance did not differ significantly between ME- and FL-derived flies in either of the two experiments (Figs. S1-2), suggesting a genetic effect primarily on LABs.

Taken together, a broad experimental approach using two different sets of wild-caught versus laboratory-reared wild flies indicates that: 1) *D. melanogaster* isolated from different geographies can select for microorganisms that confer traits consistent with their geographic adaptation; 2) the effect was more pronounced for LABs derived from natural populations than from laboratory cultures of *Drosophila;* and 3) host genetic control appeared to be stronger for LABs than for other associated bacteria.

### The microbiota influences the life history of wild D. melanogaster

The wild flies analyzed above (Fig. 3A) bore a microbiota that was consistent with the life history strategy naturally adopted by those flies (21, 34, 70), raising the question whether microbiota variation was necessary for variation in life history traits in geographically-selected fly populations. We first assessed the contribution of host genotype to the life history strategies of locally adapted fly populations by measuring the development rate and SR in ME- and FL-derived fly populations reared under microbiologically-sterile conditions. Consistent with a host genetic role, the bacteria-free FL-fly genotypes adopted a ‘fast’ strategy relative to the bacteria-free ME-flies: they were less resistant to starvation stress and completed the developmental period more quickly (Fig. 4A-B). In the final set of experiments, we assessed how the microbiota might influence the life history traits of the geographically-adapted flies (Fig. 4C-D). When reared with a single AAB- or LAB-strain that conferred extreme life history phenotypes in our previous experiments, SR and development rate in high- and low-latitude flies were more similar based on the inoculated microbes than on host genotype. These findings show that microbial influence can override genetic adaptations to differences in life history strategy.

**Figure 4.**
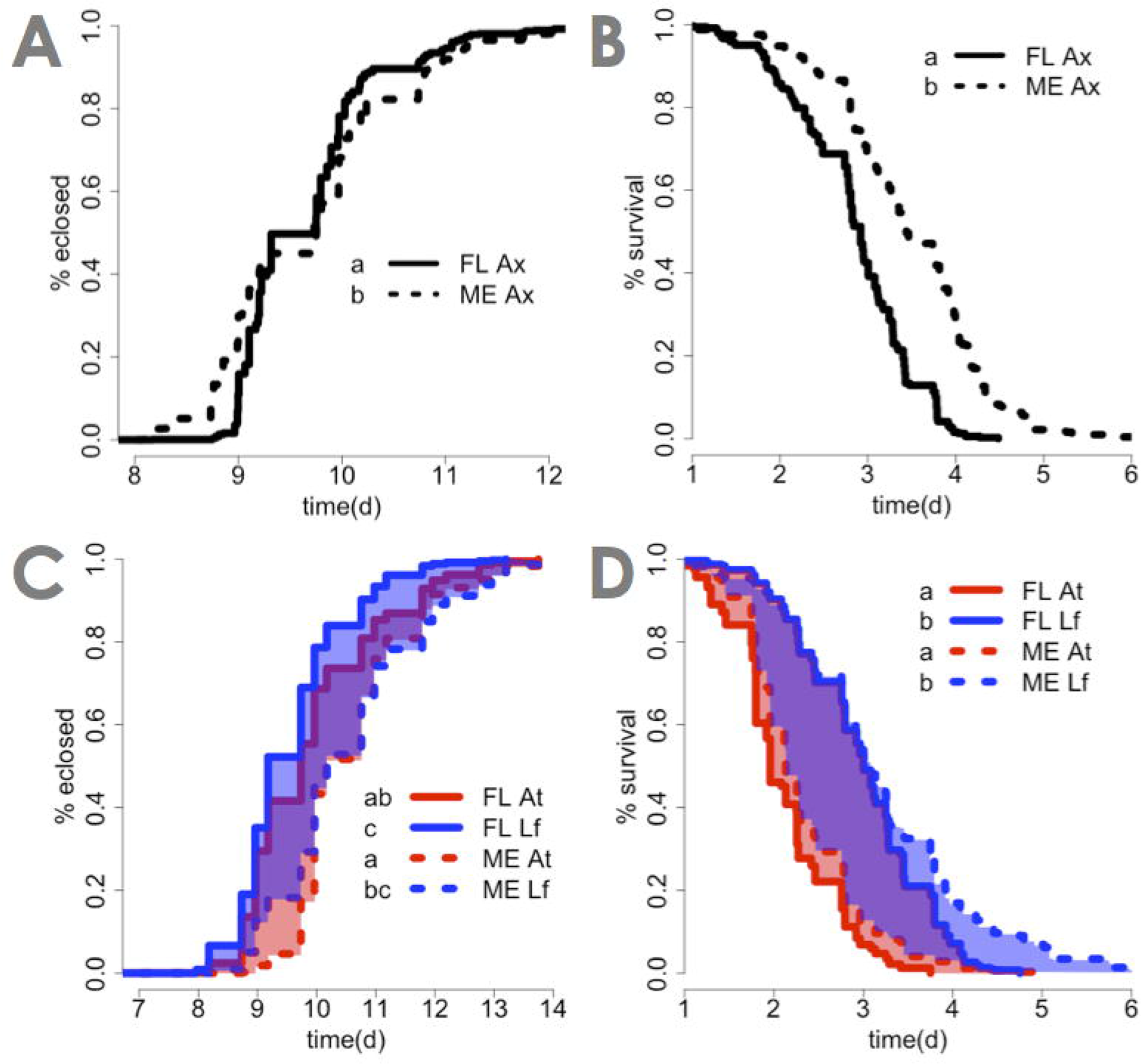
Microbial presence and identity influence life history of wild derived fly lines. Latitudinally adapted ME (dashed lines) and FL (solid lines) flies reared with different bacterial treatments were tested for variation in the period of development to eclosion (A,C) and SR (B,D) when reared bacteria-free (A, B); or in monoassociation with *A. tropicalis* (red lines) or *L. fructivorans* (blue lines) (C,D). Red or blue shading represents the capacity for variation in FL- or ME-derived flies, respectively, and shading is purple where these overlap. Except for C (triplicate vials in one experiment), all data were collected from triplicate vials in three separate experiments. Letters next to the legends represent significant differences between treatments, determined by a Cox-mixed effects survival model that included experimental replicate as a mixed effect.

## Discussion

Decades of work have established that organisms adapt to their environments in response to local environmental variation (71-75). Historically, life history adaptation has been examined from the perspective of an organism’s genetic adaptations to environmental circumstances that vary in different geographic locations, such as temperature, photoperiod, nutrient availability and predator pressure. Previous but recent work has also provided clear evidence that the microorganisms living within, on, or near a plant or animal exert substantial influence on host traits that contribute to the life history strategy, with evidence for latitudinal clines in microbiota composition in some animals, including humans, and plants (72-77). Here, we integrate these different bodies of literature to demonstrate that microorganisms associated with *D. melanogaster* are likely to influence local adaptation. The key findings of this study are twofold. First, the life history strategy of *D. melanogaster* along the “fast-slow” axis can be driven by the composition of the microbiota under experimental conditions. Second, the intrinsic host genetic factors that influence differences in life history strategy between natural populations at different latitudes include host selection of bacterial partners with congruent effects on host life history traits.

This study revealed that the bacteria can function as a rheostat to determine the fast-slow strategy adopted by the host. The molecular basis of this effect may involve bacterial production or catabolism of key metabolites. For example, bacterial production of acetic acid and other fermentation products, as well as branched-chain amino acids, can influence the activity of Insulin-like /Target of rapamycin (IIS/TOR) signaling (57, 58). IIS/TOR signaling has a central role in regulating cell and organismal growth, as well as female fecundity, and, consequently, life history strategies (78, 79). The life history traits of *D. melanogaster* **can also be influenced by** bacterial metabolism of glucose (59, 80) and methionine (64), and by B vitamin production (81, 82). The contributions of these different functions to *D. melanogaster* fitness are not known but one or more are likely to be important for *D. melanogaster* adaptation. Specifically, the insect host may utilize the composition of associated bacteria, or of bacteria in the diet, as a cue for environmental conditions (especially diet condition) in the fruit or other ephemeral habitat that support larval development. Thus, the many Acetobacteriaceae that are highly competitive in aerobic environments with high concentrations of sugars and other readily assimilated nutrients (40, 83) may represent a reliable cue for high-nutrient but ephemeral resources, favoring a “fast” phenotype of the host; while many Lactobacillales, which utilize complex carbon and nitrogen sources that are consumed more slowly (84), favor a “slow” phenotype of the host. Evidence from several studies suggest that the blend of fermentation products produced by microbial communities of different composition in the food and gut of *Drosophila* may represent a reliable cue for habitats of different nutritional content and persistence (85-87).

We have also obtained evidence that host genetic selection of its microbiota plays a key role in shaping the fast-slow strategy adopted by *D. melanogaster*. Consistent with this finding, there is a strong overlap between *D. melanogaster* genes associated with local adaptation and host-microbe interactions. For example, of 160 previously identified genes that vary in flies across latitudinal gradients (89), 45% have known or predicted (GWA, transcription) effects on microbiota interactions (44, 90, 91) relative to 31% of 13991 total *D. melanogaster* genes. (**X**^2^ = 7.03, p = 0.008). A causal role of the microbiota in these correlations is indicated by the demonstration that TAG content of *D. melanogaster* is regulated in part through genetic control of the microbiota (44). Furthermore, the genetic determinants of *Acetobacter* abundance in *D. melanogaster* include many genes that are expressed predominantly or exclusively in neurons (44), raising the possibility that sensory functions and behavioral traits (e.g. response to microbial volatiles, diet preference and feeding rate) can mediate differences in microbiota composition. These considerations raise the possibility that an individual fly might modify its lifespan/fecundity schedule in response to altered environmental circumstances, by seeking out and filtering a different suite of microorganisms. An important topic for future research is the significance of host genetic factors in driving microbiota composition in natural populations of *D. melanogaster* These host effects are more diffuse than in many associations, e.g. legume-rhizobia and *squid-Vibrio* symbioses, where exquisite specificity is dictated by defined molecular interactions (92-95) because the *D. melanogaster* association is an open system, continually exposed to microbes ingested in the food. Other deterministic factors, e.g. among-microbe interactions, as well as stochastic processes (see Introduction), may reinforce or suppress the effect of host factors on microbiota composition. Although the relative importance of these different factors is largely unknown, the demonstration of host genetic determinants of microbiota in various open associations in animals suggests that host determinants of microbiota composition contributes to the fitness of natural populations (43, 46, 47, 96-101).

This study extends our understanding of natural populations by combining studies of the microbiota in natural populations of *D. melanogaster* and laboratory analysis of precisely-controlled host-microbe combinations. For example, axenic *D. melanogaster* is a contrived state not likely experienced by flies in the wild; but was essential for establishing the influence of associated microorganisms on the flies’ genetic contributions to life history variation. Our work further emphasizes that an exclusive focus on laboratory systems can miss important interactions because it fails to reproduce the biological context underlying evolved interactions. Likely examples include the more uniform impact of *Drosophila*-derived bacteria than bacteria from other sources on fly fitness (Fig. 2A-D vs Fig. 2E-H); and, in wild flies, the difference in abundance between wild- and laboratory-fly-isolated LABs. Similarly, *Drosophila*-isolated *Acetobacter* are discordant for key functions (uric acid utilization and motility) but not taxonomy between wild-versus laboratory-flies (102). Several lines of evidence suggest that microorganisms that can persist in the flies for extended periods of time may be more prevalent in wild *Drosophila* than in long-term laboratory cultures (88, 103, 104). These issues are of general significance for the conduct of microbiome research, with parallels coming from the evidence that the microbiota of the mouse differs between laboratory inbred strains and wild (or pet-shop) mice, with associated major differences in host physiological traits, especially relating to the immune system (105-107).

The finding that the microbiota can mask host genetic determinants of life history traits is not without precedent. In particular, laboratory studies have revealed that the penetrance of various mutations on metabolic traits of *D. melanogaster* is altered, and frequently reduced, in flies colonized with microorganisms, relative to axenic flies (90). These effects cannot be attributed to a generalized dysfunction of axenic *D. melanogaster* because elimination of the microbiota has small or no fitness consequences for the host on nutrient-rich diets (55, 82), as used in this study (Fig. 2D). The comparison of life history traits in wild flies bearing either *A. tropicalis* or *L. fructivorans* showed that altering the microbial community can mask host genotype, but should not be interpreted that all AABs or LABs confer life history traits of the same magnitude. Numerous studies of traits in monoassociated *Drosophila* reinforce the expectation that the microbial influences are species-specific and possibly even strain-specific (58-60, 64, 67, 102, 108-111). Further research is required to elucidate the underlying mechanisms, and to establish the extent to which the impact of individual microbial taxa on host life history traits in monoassociations (as used in these experiments), or controlled multi-species associations (112) may be displayed in the taxonomically diverse microbial communities in natural fly populations.

In conclusion, the impact of the microbiota on the life history strategy of *D. melanogaster* is most unlikely to be a unique trait of this insect species. Microbial influence on life history traits of animals may be widespread, representing an important, but hitherto neglected, determinant of intraspecific variation, including local adaptation. We recommend that analysis of the microbiota is included as an integral part of research on life history traits and local adaptation in animals, to determine the magnitude of microbial effects in different systems, and to establish the proximate and ultimate mechanisms.

## Materials and Methods

### Fly rearing and bacterial culture conditions

Standard fly rearing conditions were at 25°C using a 12 hour light-dark cycle on a yeast-glucose (YG) diet (10% yeast, 10% glucose, 1.2% agar) containing 0.042% propionic acid and 0.04% phosphoric acid (67)). Fly lines are listed in Table S3. *Wolbachia* status of the flies was determined using the wsp691-R (5’-AAAAATTAAACGCTACTCCA-3’) and wsp81-F (5’-TGGTCCAATAAGTGATGAAGAAAC-3’) as described previously (113).

To control bacterial exposure to particular microbial partners and to test for the influence of individual microbes on life history traits, we reared flies under bacteria-free conditions or from bacteria-free eggs with an inoculated, defined microbiota. Fly eggs were collected from grape juice plates, dechorionated in 0.6% sodium hypochlorite for two 2.5 minutes washes, rinsed three times with sterile water, and transferred to sterile YG diet (no acid preservative added) in a biosafety cabinet, as in our previous work (54). Bacteria-free eggs were left unmanipulated, or, to rear flies with a defined microbiota, were inoculated with 50 μl bacterial culture that had been grown overnight and normalized to OD_600_=0.1.

Bacterial strains (Table S4) were cultured on specific media: modified MRS medium (mMRS; 1.25% peptone, 0.75% yeast extract, 2% glucose, 0.5% sodium acetate, 0.2% dipotassium hydrogen phosphate, 0.2% triammonium citrate, 0.02% magnesium sulfate heptahydrate, 0.005% manganese sulfate tetrahydrate, 1.2% agar (67)), potato medium (pot; [Sigma P6685]), lysogeny broth (LB; 1% tryptone, 0.5% yeast extract, 0.5% sodium chloride), and brain heart infusion (BHI, Sigma 53286). All strains were grown at 30°C except *Escherichia coli*, which was grown at 37°C. Strains grown under normoxia were shaken (liquid) or under ambient laboratory conditions (solid). Strains requiring hypoxia were grown statically (liquid) or in a sealed container flooded with CO_2_ (solid).

### Bacterial abundance

Bacterial abundance was assessed in whole body fly homogenates. Flies from each vial were anesthetized and a pool of five flies was homogenized in 125 μl homogenization buffer (10 mM Tris, pH 8, 1 mM EDTA, 0.1% Triton X-100 as in (59)) with 125 μl Lysing Matrix D ceramic beads (MP Biomedicals 116540434) by shaking for 30-60s at 4.0 m/s in a FastPrep-24 or 1500 RPM for 2 min on a GenoGrinder 2010. The homogenate was plated onto mMRS medium twice, with dilution plating under normoxic or hypoxic conditions for enumeration of bacterial abundance; and a spot-test under the reverse (hypoxia or normoxia) conditions to test for contamination. After incubation the colony morphologies were inspected visually to confirm strain identity. Where ≥ 200 CFU fly^-1^ of the expected bacterial strain were detected, the strain was deemed ‘present’. Differences between *Acetobacter* strains could usually not be determined by colony morphology, so *Acetobacter* contamination of other *Acetobacter* strains cannot be ruled out.

### Development rate of the flies

*D. melanogaster* development rate to pupariation or eclosion was determined by counting the number of pupae formed at 1, 6.5, and 10 hours into the daily light cycle. For Fig. 4A, three separate experiments each with triplicate fly vials were performed for each treatment. For Fig. 4C, eclosion was measured and analyzed in one experiment (triplicate vials).

### Starvation resistance (SR)

SR was determined in pools of ten 5-7-day-old female flies. Female flies were separated from under light CO_2_-anesthesia, and then incubated in fly vials containing 5 ml 1% agarose under standard fly rearing conditions. Fly mortality was recorded every six hours until all flies in a vial were dead. Three separate experiments each with triplicate fly vials were performed for each treatment.

### Glucose content

Glucose content was measured from homogenized pools of five 5-7 day-old female flies as in our previous work (59). Briefly, the pool of flies was homogenized in 10 mM Tris – 1 mM EDTA – 0.1% Triton X-100 buffer and analyzed by the Sigma Glucose Assay kit (GAGO20-1KT) according to manufacturer instructions.

### Lifespan

*D. melanogaster* adult lifespan was measured by recording the number and sex of dead flies and transferring surviving flies to fresh sterile diet every 2-3 days until all flies in a vial were dead. For every transfer of adult flies (P generation) to fresh diet, one spent vial per week was retained for 2-3 weeks, when the offspring (F1 generation) were homogenized to check for bacterial persistence and contamination during transfer. Where ≥ 200 CFU fly^-1^ of an unexpected bacterial species were detected in two consecutive weeks, the flies were deemed contaminated from the first date contamination was detected. For survival analysis, flies were marked as leaving the experiment alive at that time; for fitness analyses the vials were discarded. At least three separate experiments with triplicate fly vials were performed for each treatment.

### Fecundity

*D. melanogaster* fecundity was determined over a 35 day period in monoassociated flies and defined as the number of F1 offspring that reached pupation in twice-weekly measures over a 4-week interval. First, 30-60 P generation *D. melanogaster* per vial were monoassociated with different bacterial strains. After >90% of flies had eclosed, P generation adults were serially transferred to sterile YG diet twice per week, between 8 and 10 hours into the daily light cycle, for 4 weeks. At 18 hours later (between 2 and 4 hours into the daily light cycle the following day), the flies were transferred to new, sterile food until the next cycle of 18 hour-fecundity measures. To ensure that the flies did not lose their bacteria during the frequent transfers, the diets for fecundity measures were inoculated with 50 μl OD_600_=0.1 normalized bacteria two days before flies were transferred. Bacteria were normalized in PBS to nutrient supplementation from spent media during the transfer. Spent vials of serially-transferred P generation flies were stored at 20°C on an uncontrolled light cycle, approximately 6AM – 6PM, and the number of pupae that formed after ~ two weeks was calculated and normalized to the number of P generation females per vial and the exact time the P generation spent in the vial. Contamination during transfers was monitored weekly as in the lifespan analysis. If at least two consecutive F1 vials contained ≥200 CFU fly^-1^ of an unexpected bacterial colony morphology, fecundity data for the P flies was discarded from all analyses. Early fecundity values (12-15 days post egg deposition) were used for the correlation analysis. For the other analyses, differences in fecundity between experiments were calculated on a 2-4 day sliding scale. Three separate experiments with triplicate vials were performed on three consecutive days and data were pooled and analyzed based on date of vial transfers.

Fitness was calculated from the twice-weekly fecundity data paired with fly survival measurements over the same period. Leslie matrices were created (68, 69) and eigenvalue lambda was calculated in R using the eigen function for each replicate vial. For each of fecundity and fitness measures, a Kruskal-Wallis rank sum test was used to assess treatment-level differences. If the Kruskal-Walis test was significant, then differences between treatments were assessed by a mixed model (114) and a Tukey post-hoc test (115) for multiple comparisons‥

### 16S rRNA marker gene analysis

To test for microbiota composition of wild *D. melanogaster* the V4 region of the 16S rRNA marker gene was amplified from whole body homogenates of wild flies. In fall of 2009, wild flies were collected to empty fly vials, and *D. melanogaster* were sorted from the mixed species pools (if any) within 16 hours and stored in ethanol with < 100 other flies from the sample collection. DNA was extracted from triplicate pools of 5 whole-body flies by a salting out procedure (116). Briefly, fly bodies were homogenized in enzymatic lysis buffer with 20mg/ml lysozyme (Amresco, 0663) and disrupted using glass beads. Cells were then lysed via incubation with 10X extraction buffer and proteinase K. After incubation with 3 M sodium acetate, DNA was extracted from the pellet using 100% isopropanol, rinsed in 70% ethanol and resuspended in sterile TE buffer. From these extracts, the V4 region of the 16S rRNA gene was amplified as described previously (117). Sequences were normalized using the Invitrogen Normalization kit and sequenced via 2 x 250 Illumina v2 chemistry on a HiSeq 2500 at the BYU DNA sequencing center. Sequence reads are available at NCBI (*Accession number forthcoming*).

Operational taxonomic units were clustered and assigned to the sequencing data in QIIME 1.9.1 using UCLUST with open-reference OTU picking and the GreenGenes Core reference alignment at 97% similarity (118-122). Taxonomy was assigned with the RDP Classifier 2.2 (122) and the GreenGenes reference database (123, 124). *Wolbachia* reads were filtered from the OTU table, which was rarefied to 65 reads per sample, which was still sufficient to near-saturate most samples (Fig. S3). Spearman rank correlations between order-level OTU classifications (LABs = Lactobacillales, AABs = Rhodospirillales, almost all from Acetobacteraceae), were performed in R.

### Host selection on the microbiota

To test for host genetic selection on microbiota composition of clinally-adapted fly lines, we enumerated bacterial abundance in *D. melanogaster* administered defined bacterial taxa. The defined 5-species bacterial community by inoculated bleach-sterilized fly eggs with an equal-ratios mixture (normalized to OD_600_=0.1) of *A. tropicalis, A. pomorum, L. brevis, L. fructivorans*, and *L. plantarum* (Table S4, 5-sp). The microbial communities associated with the flies were assessed by dilution plating whole body homogenates of 5 pooled male flies. Whole-body fly homogenates were prepared in 125 μl TET buffer (10 mM Tris pH 8, 1 mM EDTA 0.1% Triton X100) with an equal volume of Lysing Matrix D ceramic beads (MP Biomedicals) by shaking on a FastPrep24 at 4.0 m/s. Visual differences in colony color and morphology were used to distinguish *Lactobacillaceae* (large, white or yellow) and *Acetobacter* (small, tan) colonies. In a followup study the same fly lines were reared with a 6-species inoculum composed of 4 *Lactobacillaceae* species isolated from wild Drosophila and the two previously used *Acetobacter* strains, which were isolated from laboratory *Drosophila* (strains DmW_98, DmW_103, DmW_181, DmW_196, apoc, atrc; Table S4). Samples were prepared and microbiota composition was enumerated by the same methods as above.

### Statistical analyses

To define the relationships between life history traits in monoassociated CantonS flies, Pearson (if normal by a Shapiro test) or Spearman (if not normal by a Shapiro test) rank correlations between mean phenotype values were calculated in R (125). All assays were performed on the same fly genotype using the same diet formulation, bacterial strains, and general methods. One major bifurcation in the data is that the previously published TAG content, development rate, feeding rate, and glucose content data were collected in Ithaca, NY; whereas the SR, lifespan, and fecundity data were collected in Provo, UT. The previously published data were used for different purposes, and this is a nonredundant analysis of those data.

To test for genotype by microbiota differences in development rate or SR, Cox mixed effects models were used (126, 127). Model complexity was determined by selecting the model with the lowest AIC score. Differences in microbiota composition were defined by Kruskal-Wallis rank sum tests or linear mixed effects models with a binomial family (114).

To test for statistically significant differences in the CFU abundance data we employed 2 approaches. First, we examined the raw CFU counts as a ratio of *Lactobacillaceae* to *Acetobacter* abundance in flies grouped by geographic cline or fly genotype using a generalized linear mixed effects model with a binomial family (114, 115, 125). If there was a cline-specific difference, then the difference in the microbiota of fly lines was tested using fly genotype as a fixed effect instead of geographic cline. We also compared the absolute abundances of the bacteria. A Shapiro test was used to confirm the CFU abundances were not distributed normally and differences between bacterial abundances in lines from ME- and FL-geographies were determined by a Kruskal-Wallis test. Differences in bacterial abundances by fly line were determined by a Dunn test with Benjamini-Hochberg correction after confirming a significant line effect by a Kruskal-Wallis test. All tests were performed in R. (125, 128, 129).

### Data sharing

All fly lines and bacterial strains will be made freely available upon request to the corresponding author. Sequence data are uploaded to public databases (*Accession number forthcoming*).

## Acknowledgements

We thank Drs. Jerry Johnson and Byron Adams for helpful discussions.

## Supporting information captions

Figure S1. A,B) Relative CFU abundances of *Acetobacter* (red) and *Lactobacillus* (blue) in gnotobiotic 5-species fly lines isolated from wild-caught *D. melanogaster* in ME or FL, USA. Differences between treatments were determined by a generalized linear mixed effects model with a binomial family followed by ANOVA and, where p< 0.05 for the fixed effect, post-hoc Tukey tests. Absolute CFU abundances of *Acetobacter* (C,D) and *Lactobacillus* (E,F) in gnotobiotic 5-species fly lines derived from wild-caught *D. melanogaster* in ME, USA (blue) and FL, USA (red). All the inoculated *Lactobacillus* species were isolated from laboratory-reared *D. melanogaster* (67). Differences between treatments were determined by a Kruskal-Wallis test which, if p < 0.05, was followed by a Dunn’s test.

Figure S2. A, B) Relative CFU abundances of *Acetobacter* (red) and *Lactobacillaceae* (blue) in gnotobiotic 6-species (C,D) fly lines isolated from wild-caught *D. melanogaster* in ME or FL, USA. Differences between treatments were determined by a generalized linear mixed effects model with a binomial family followed by ANOVA and, where p< 0.05 for the fixed effect, post-hoc Tukey tests. Absolute CFU abundances of *Acetobacter* (C,D) and *Lactobacillaceae* (E,F) in gnotobiotic 6-species fly lines derived from wild-caught *D. melanogaster* in ME, USA (blue) and FL, USA (red). All the inoculated *Lactobacillus* species were isolated from wild *D. melanogaster* (130-132); data not shown}. Differences between treatments were determined by a Kruskal-Wallis test which, if p < 0.05, was followed by a Dunn’s test.

Figure S3 Rarefaction analysis of microbiota data in Fig. 3A, performed using QIIME 1.9.1.

